# Host genetics and pre-vaccination blood transcriptome as determinants of vaccine-induced immunity to Influenza A virus in swine

**DOI:** 10.64898/2026.02.06.700797

**Authors:** Fany Blanc, Tatiana Maroilley, Gaëtan Lemonnier, Laure Ravon, Yvon Billon, Olivier Bouchez, Marie-Hélène Pinard-van der Laan, Jordi Estellé, Claire Rogel-Gaillard

**Affiliations:** Université Paris Saclay, INRAE, AgroParisTech, GABI, 78350 Jouy en Josas, France; GenESI, INRAE, 17700 Surgères, France; INRAE, GeT-PlaGe, Genotoul, 31326 Castanet-Tolosan, France

**Keywords:** vaccination, prediction, GWAS, blood transcriptome, biomarker, antibody response, Influenza A virus, pig

## Abstract

Influenza A virus (IAV) is a major respiratory pathogen in pigs, causing diseases that have significant economic and potential public health consequences. Vaccine effectiveness varies among animals, impacting long-term herd protection due to individual variabilities in antibody levels and persistence over time. Our aim was to identify the genetic factors and pre-vaccination blood transcriptomic profiles that influence immune response levels to the IAV vaccine. A total of 187 piglets were vaccinated at weaning (28 days of age, 0 days post-vaccination, dpv) and boosted three weeks later, and humoral responses were assessed until slaughter (21, 28, 35, and 118 dpv) by measuring serum IAV-specific IgG and hemagglutination inhibition (HAI) titers.

The results revealed varying antibody responses and persistence. Genome-wide association studies identified two loci on chromosomes SSC5 and SSC8 associated with a persistence of HAI titers until slaughter. Pre-vaccination blood transcriptomic analyses showed that early and post-boost antibody responses (21, 28 and 35 dpv) and long-term persistence (118 dpv) were associated with distinct baseline immune programs, with extracellular matrix and myeloid-related signatures predicting strong early and peak responses, whereas interferon-related signatures were linked to reduced long-term antibody persistence. Our results highlight the importance of considering the role of immune competence and genetics in vaccine responses in pigs and suggest candidate biomarkers to improve vaccination strategies within breeding programs.

## Introduction

Influenza A virus (IAV) is the causative agent of flu, a highly contagious respiratory disease that can lead to significant illness and death in both humans and other animal species, including poultry and swine. In pigs, this disease contributes to the porcine respiratory disease complex, resulting in a significant economic impact because of reproductive failure in sows and weight loss in growing pigs ^1,2^. Pigs are often considered as “mixing vessels” or transmission agents in the adaptation of avian influenza viruses to mammals, including humans ^3,4^, raising concerns that these viruses may cause zoonotic infections. This zoonotic risk underscores the importance of controlling IAV in pigs, not only for economic and animal welfare reasons, but also from a One Health perspective that integrates animal and public health.

Vaccination remains the most effective strategy to control and prevent infectious diseases in livestock, including IAV infection in pigs ^5^. Ideally, a vaccine should be cost-effective, easy to administer, safe, and capable of inducing a strong and long-lasting immune response. Under field conditions, pigs may already harbor some level of pre-existing immunity, either through maternally derived antibodies or as a result of previous infections. Such immunity may be partial and transient and does not necessarily prevent infection or transmission ^6^. Therefore, as with other vaccines, IAV vaccines must even be effective in the presence of such immunity. Improving vaccine efficacy is therefore crucial to achieving stronger and more durable protection, as well as limiting virus circulation at the herd level.

Considerable inter-individual variability in antibody responses to vaccination has been reported not only for IAV ^7^, but also for other vaccines in pigs, such as those against porcine reproductive and respiratory syndrome virus ^8^, tetanus ^9^, bacterial antigens ^10^, *Mycoplasma hyopneumoniae* ^11–13^. Understanding and predicting this variability is essential in order to optimize vaccination strategies against IAV and other infectious agents.

Numerous studies in humans have investigated the genetic basis of inter-individual variability in vaccine responses. Single-nucleotide polymorphisms (SNPs) in immune-related genes have been found to be associated with differences in immune responses to vaccines against measles and rubella ^14,15^, smallpox ^16^, capsular group C meningococcal, *Haemophilus influenzae* type b, tetanus toxoid ^17^ or *Bacillus anthracis* ^18^. Furthermore, meta-analyses and studies evaluating multiple vaccines in the same population have demonstrated that specific HLA class II alleles modulate antibody responses to several human vaccines ^19,20^, suggesting that genetic determinants of vaccine responsiveness may act through shared immune pathways rather than being vaccine-specific. More recently, similar genetic influences have also been reported with respect to SARS-CoV-2 vaccination ^21,22^. In pigs, similar findings have been reported by our group and others, with genomic regions associated with vaccine responses against porcine reproductive and respiratory syndrome virus ^8,23^, classical swine fever ^24,25^ and *Mycoplasma hyopneumoniae* ^13^, confirming that host genetics influence the magnitude of vaccine responses.

In humans, the transcriptomic profiling of immune responses following IAV vaccination has identified blood gene expression signatures and co-expression modules that correlate with, and even predict, the magnitude of antibody responses ^26–30^. These studies demonstrated that blood transcriptomics offer a powerful tool to evaluate vaccine-induced immune responses and elucidate the mechanisms underlying protective immunity. For example, the early induction after vaccination of type I interferon (IFN-I)–driven innate immune pathways has been associated with stronger antibody responses. More recently, single-cell transcriptomic analyses of porcine immune cells provided new insights into B- and T-cell clonal expansion and gene regulatory signatures associated with vaccination and infection ^31,32^.

Moreover, baseline (pre-vaccination) transcriptomic profiling has revealed gene expression patterns and immune modules that predict subsequent antibody responses to IAV vaccination ^29,33^. Thus, pre-existing immune states, and not just early innate responses, contribute to vaccine responsiveness in humans. Comparable approaches have recently been applied in pigs. Pre-vaccination blood transcriptomic profiling has been shown to identify gene signatures and functional modules that correlate with, and predict, the antibody response to *Mycoplasma hyopneumoniae* vaccination ^13^, thus confirming that baseline immune transcriptional activity, as observed in humans, also influences vaccine responsiveness in pigs. However, the mechanisms underlying such variability remain poorly characterized for other porcine vaccines, and particularly those targeting IAV.

Building on the results for other porcine pathogens, we hypothesized that both host genetic variation and pre-vaccination blood transcriptional activity contribute to the inter-individual variability observed in antibody responses to IAV vaccination in pigs. To test this hypothesis, we combined genome-wide association studies (GWAS) and pre-vaccination blood transcriptomic profiling in the same experimental cohort, according to the same experimental design as that used to analyze individual variability in the response to *Mycoplasma hyopneumoniae* vaccine ^13^. This integrative approach was designed to identify genetic loci and baseline transcriptional signatures associated with vaccine-induced antibody responses, to provide clues on underlying biological mechanisms and to suggest candidate biomarkers of vaccine efficacy in pigs.

## Results

### Cohort overview and sampling

A total of 187 piglets were vaccinated at weaning (around 28 days of age, 0 days post-vaccination, dpv) and received a booster vaccination three weeks later (21 dpv). Sixty-four non-vaccinated littermates served as controls. All animals were monitored from birth until slaughter, which occurred at approximately 146 days of age. Humoral responses were assessed at 21, 28, 35, and 118 dpv by measuring serum IAV-specific IgG and hemagglutination inhibition (HAI) titers. These time points captured the early immune response (21 dpv), the post-boost peak response (28 and 35 dpv), and the persistence of the response (118 dpv) (Fig. 1a).

**Figure 1.**
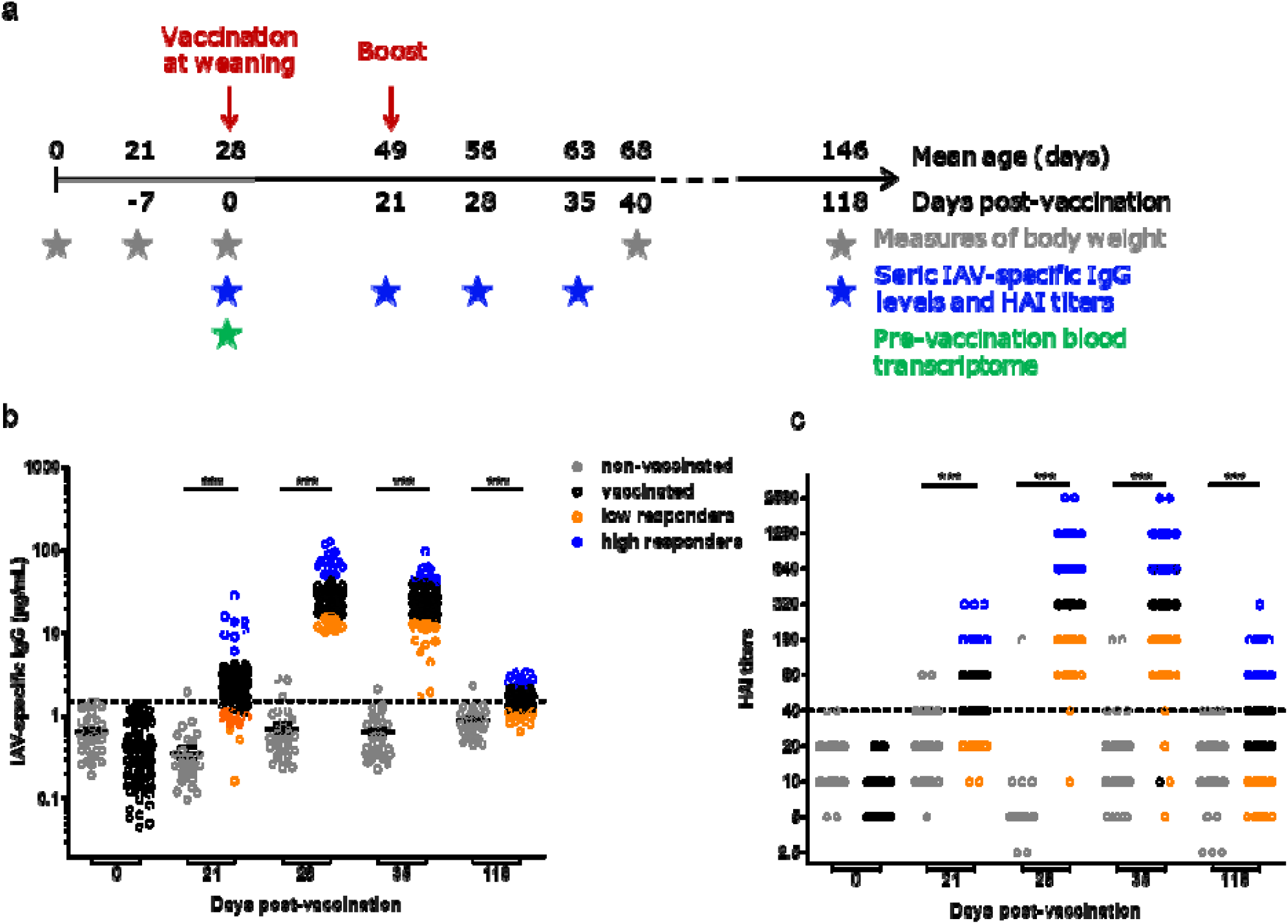
IAV antibody response in pig sera after vaccination. (a) Experimental design showing the vaccination schedule, body weight measurements, and blood sampling protocol. Grey stars indicate body weight measurements, blue stars point to the collection of blood samples to monitor antibody responses to vaccination, and the green star indicates blood sampling for pre-vaccination blood transcriptome analysis. (b) Serum levels of IAV-specific IgG measured at 0, 21, 28, 35, and 118 days post-vaccination (dpv) in vaccinated and non-vaccinated animals from litters without maternally transmitted IAV-specific IgG on the day of vaccination (groups V without_ MDA and NV without MDA, respectively). (c) HAI titers in sera from vaccinated and non-vaccinated animals at 0, 21, 28, 35, and 118 dpv (all piglets included). The dotted line indicates the seroprotection threshold defined as HAI ≥40. Statistical analysis was performed using unpaired *t*-tests between control and vaccinated animals (***p <0.001).

### Baseline presence of maternally derived antibodies (MDA) in some animals and analysis sets

At baseline on the day of vaccination (0 dpv), some piglets already exhibited IAV-specific IgG, indicating the presence of maternally derived antibodies (MDA). Based on IgG levels at 0 dpv, piglets were subsequently categorized according to their MDA status as follows: MDA□: all littermates with MDA <1.5 µg/mL (98 vaccinated and 34 control pigs, from 26 litters); MDA□: ≥50% of littermates with MDA >1.5 µg/mL (70 vaccinated and 23 control piglets, from 16 litters); and non-classified (NC), which did not meet the criteria (10 vaccinated and 7 control piglets; see Supplementary Table 1).

Thus, baseline IAV-specific IgG levels in MDA□ piglets were lower than in MDA piglets (Supplementary Fig. 1a). Consistent with this observation, sows from the MDA□ group displayed a broad range of serum IAV-specific IgG levels (4.3–772 µg/mL), while sows from the MDA group harbored lower IgG levels (0.3–15.4 µg/mL) (Supplementary Fig. 1b).

Because maternally derived IAV-specific IgG was indistinguishable from vaccine-induced IgG in MDA piglets (Supplementary Fig. 1c), subsequent IAV-specific IgG analyses were limited to the MDA□ group. In contrast, HAI titers were analyzed in the entire cohort, as all MDA□ piglets exhibited HAI <40 at 0 dpv (Supplementary Fig. 2a), and only two sows (one from each MDA group) showed HAI titers ≥40 (Supplementary Fig. 2b).

### Individual variability of humoral responses and identification of extreme responders

#### IAV-specific IgG kinetics (MDA^-^ group)

Control piglets maintained IAV-specific IgG levels lower than 1.5 µg/mL after 0 dpv, with only six occasional values slightly exceeding this threshold. In contrast, vaccinated piglets showed a significant vaccine-induced humoral response. At 21 dpv, most piglets (76%) had seroconverted and showed IgG levels above the positivity threshold. Overall antibody levels remained relatively low at this early time point but already exhibited considerable variability among individuals (mean 3.25 ± 3.77 µg/mL; CV = 115.9%). A substantial rise in IAV-specific IgG levels was observed at 28 dpv, one week after the booster dose, with all animals displaying levels well above the threshold (33.41 ± 23.25 µg/mL; CV = 69.6%). This strong response persisted at 35 dpv (27.73 ± 14.73 µg/mL; CV = 53.3%) and was considered as the maximum response. Thereafter, antibody levels declined, and by 118 dpv they had largely returned to low levels (1.68 ± 0.61 µg/mL; CV = 36.3%). Despite this decline, levels in 56% of the piglets remained above the predefined positivity threshold, and the antibody levels of vaccinated animals were still higher than those of the control group (Fig. 1b; Supplementary Table 2). No significant effects related to sex, litter or batch were detected, although there were weak trends for age at weaning at 21 dpv and batch at 118 dpv (p < 0.1; Supplementary Table 3).

Extreme responders were defined using log_10_-transformed IAV-specific IgG values: high responders were those with values above the mean + 1 SD, and low responders were those with values below the mean - 1 SD. The resulting count of extreme responders was: 8 high/14 low (21 dpv), 18 high/16 low (28 dpv), 12 high/17 low (35 dpv), and 14 low/14 high (118 dpv) (Fig. 1b; Supplementary Table 2).

#### HAI kinetics (full cohort)

Seroprotection rates (HAI ≥40) rose from 85.5% at 21 dpv (with a median of 80) to 99.5% at 28 dpv and 98.4% at 35 dpv (with medians of 320) and remained at 47.9% before slaughter at 118 dpv (with a median of 20) (Fig. 1c). MDA status had no effect on HAI titers at any time points (Supplementary Fig. 2c; Supplementary Table 4). Batch effects were observed at 21, 35, and 118 dpv, while sex effects were noted at 28 dpv, with females exhibiting higher HAI titers than males (*p* <0.05; Supplementary Table 3).

Extreme responders were defined based on the following HAI titer thresholds: 21 dpv: low ≤ 20, high ≥ 160; 28 and 35 dpv: low ≤ 160, high ≥ 640; 118 dpv: low ≤ 10, high ≥ 80. These thresholds yielded 21 high/26 low (21 dpv), 68 high/44 low (28 dpv), 52 high/71 low (35 dpv), and 30 high/54 low (118 dpv) (Fig. 1c).

### Consistency between IAV-specific IgG and HAI

Within each assay, the strongest correlations were observed between successive sampling points. Simple Pearson correlations revealed the highest correlations between antibody responses at the closest time points, with IAV-specific IgG showing a correlation of r = 0.81 between 28 and 35 dpv, and HAI titers showing a correlation of r = 0.68 over the same interval. Correlations decreased as the time lapse between sampling points increased and were found to be the lowest for comparisons with levels at 118 dpv (Fig. 2). To confirm these results, we also applied a statistical model that accounted for the fact that the same animals were sampled repeatedly over time. This model (mixed-effects with an autoregressive structure) similarly showed that measurements taken closer together in time were more strongly correlated, supporting the Pearson correlation results (Supplementary Table 5).

**Figure 2.**
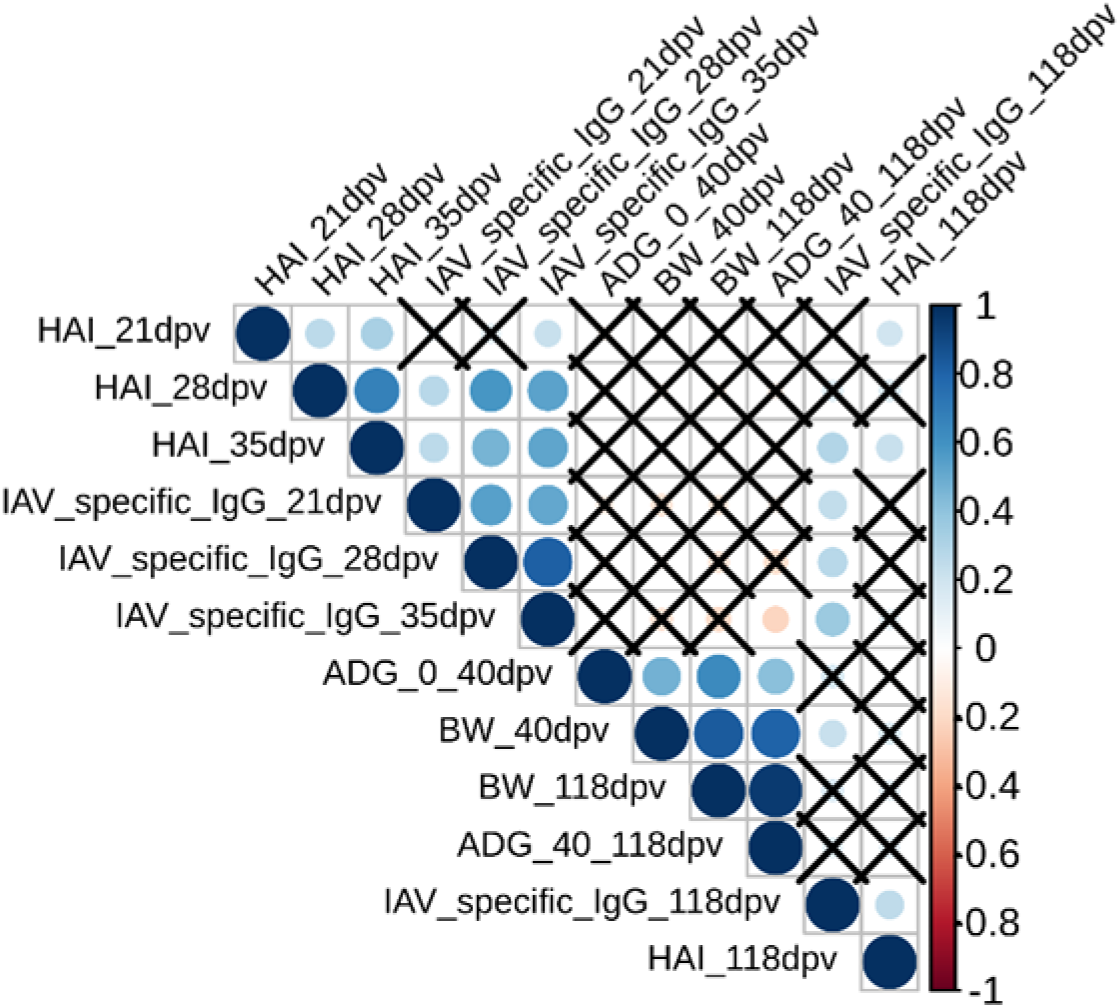
Correlations between antibody levels and growth traits after vaccination against IAV. Correlation matrix showing Pearson correlation coefficients between serum IAV-specific IgG levels and HAI titers measured at 21, 28, 35, and 118 days post-vaccination (dpv); body weights (BW) at 40 and 118 dpv; and average daily gain (ADG) from 0–40 dpv and 40–118 dpv. Non-significant correlations (*p* >0.05) are crossed out.

Across assays, significant positive correlations were found overall (Fig. 2). Notably, HAI at 28 and 35 dpv correlated with IAV-specific IgG at 21, 28 and 35 dpv, with the highest observed correlation (r = 0.58) between assays at 28 dpv. Additionally, HAI at 35 dpv correlated with IAV-specific IgG at 118 dpv (r = 0.29). These findings highlight the complementary roles and information provided by IgG levels and HAI titers in characterizing vaccine responses.

### Pig growth and specific IAV-IgG antibody response: evidence of a slight trade-off

Body weights (BW) at 40 dpv and 118 dpv were strongly correlated (r = 0.83), whereas average daily gain (ADG) during the post-weaning and growing periods were moderately correlated (r = 0.42, Fig. 2). Two significant correlations were detected between growth and immune traits: IAV-specific IgG levels at 35 dpv were negatively correlated with ADG during the growing period (r = – 0.21), while IgG levels at 118 dpv were positively correlated with BW at 40 dpv (r = 0.23).

High responders at 35 dpv (based on either IAV-specific IgG or HAI titers) had lower BW at 118 dpv, showing a decrease of 7.3% and 4.2%, respectively (*p* = 0.03 and 0.021; see Supplementary Table 6). For high HAI responders at 35 dpv, a significant 5.0% reduction in ADG during the growing period was also observed (*p* = 0.021, Supplementary Table 6). Similar but non-significant trends were observed for vaccine responses at 28 dpv. Overall, these results indicate a slight trade-off between growth performance and antibody response intensity, as revealed by either specific IgG levels or HAI titers.

### GWAS: identification of two loci associated with a persistence of HAI titers

To explore the genetic architecture of antibody responses to the IAV vaccine, 187 vaccinated pigs were genotyped using the 658K Axiom Porcine Genotyping Array (Affymetrix). Genome-wide association studies (GWAS) were conducted on 425,689 high-quality SNPs to identify loci associated with HAI titers at different time points. Significant associations (FDR <0.05) were only detected for HAI titers at 118 dpv (Fig. 3; Supplementary Table 7), with two regions identified: SSC5 (11,296,931–11,302,721) and SSC8 (20,193,741–20,258,950).

**Figure 3.**
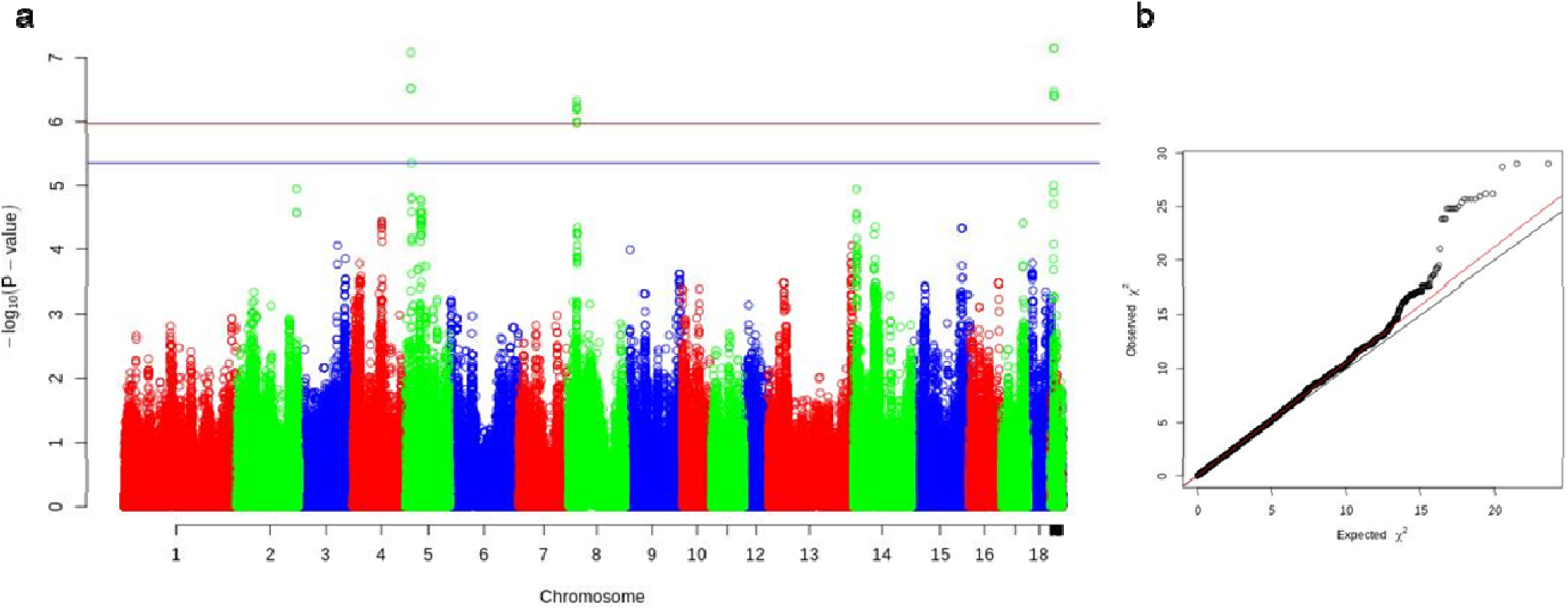
Genome-wide association analysis for HAI titers at 118 days post-vaccination (dpv). (a) Manhattan plot showing –log (*p*) values from the genome-wide association study (GWAS) and imputation analysis plotted against chromosomal positions based on the Sscrofa11.1 swine reference genome assembly. The blue line indicates the suggestive association threshold (FDR = 0.1), and the red line indicates the genome-wide significance threshold (FDR = 0.05). (b) Quantile–quantile (QQ) plot showing the expected versus observed distributions of test statistics across single-nucleotide polymorphisms (SNPs).

Within the extended SSC5 interval (QTL ±1 Mb), several genes with strong relevance to humoral immunity were identified. The strongest candidates included *IL2RB, CSF2RB, RAC2* and *CARD10*, all involved in T-cell help, B-cell activation or NF-κB-dependent survival pathways supporting long-lived antibody production. Additional genes implicated in innate immune regulation (*NCF4, C1QTNF6*), stromal remodeling (*TIMP3*) and redox homeostasis (*TMPRSS6, TXN2*) also emerged as plausible contributors.

For the SSC8 region, two genes were localized within the restricted QTL interval defined by the significantly associated SNPs (*CCKAR* and *TBC1D19*). Extending the window by ±1 Mb revealed additional biologically relevant immune genes; notably *RBPJ*, a central regulator of the Notch pathway, and *STIM2*, which regulates sustained Ca² signaling that is critical for long-term lymphocyte activation and plasma cell persistence. These genes represented strong candidates underlying variations in antibody persistence.

Only MDA animals (n = 98) could be analyzed for IAV-specific IgG levels, and no significant associations were detected, likely reflecting limited statistical power in this subset.

### Analysis of pre-vaccination blood transcriptomes reveals biological links with the magnitude of the antibody response to the IAV vaccine

#### Differential gene expression between high and low antibody responders before vaccination

Pre-vaccination blood transcriptomes (0 dpv) were generated from a subset of 91 vaccinated pigs without MDA. Differential expression analyses were conducted to compare animals classified as high or low responders to IAV vaccination at 21, 28, 35, and 118 dpv, considering both IAV-specific IgG levels and HAI titers. The numbers and lists of significantly differentially expressed (DE) genes (FDR <0.05) for both immune traits across all time points are provided in Table 1 and Supplementary Table 9.

**Table 1:**
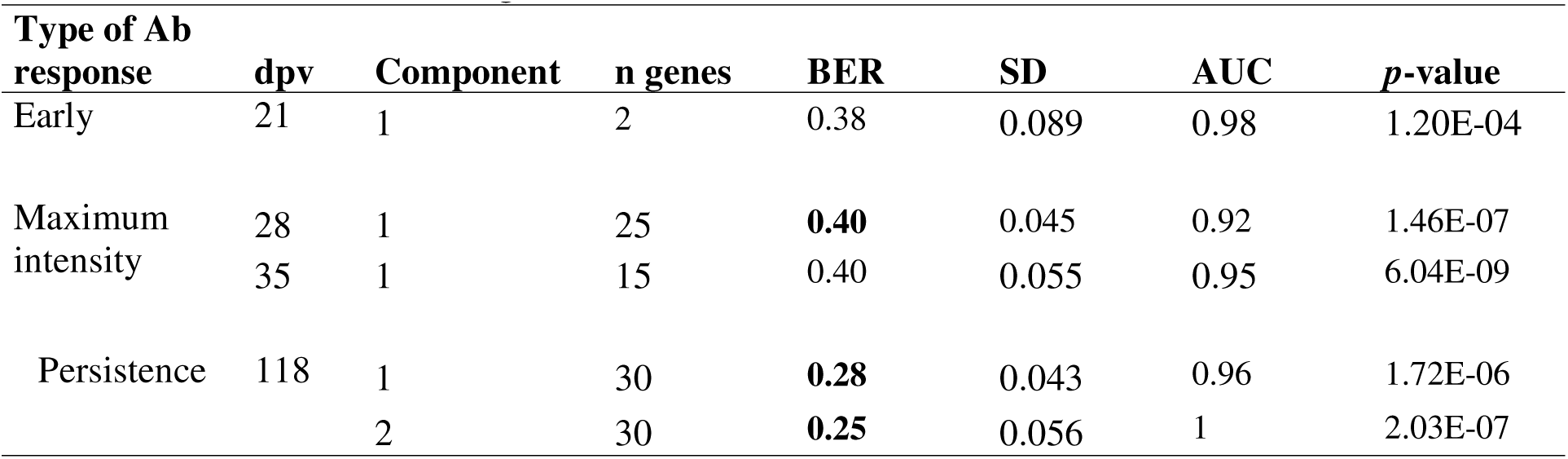
Differentially expressed genes between extreme Ab responders to IAV vaccination at the various days post-vaccination (dpv). BER: Balanced Error Rate (significant BER are in bold). AUC: Area Under the Curve.

Only a few DE genes distinguished high and low responder groups at the peak of Ab production (28 and 35 dpv) for both IAV-specific IgG or HAI titers. By contrast, 65 DE genes were detected among extreme early responders (based on IAV-specific IgG at 21 dpv), and 243 DE genes were identified among animals displaying divergent Ab persistence profiles (based on HAI titers at 118 dpv).

#### Functional analysis of pre-vaccination blood transcriptomic profiles among high and low responders at 21 dpv

Functional enrichment analyses were performed on annotated DE genes using Ingenuity Pathway Analysis (IPA) to identify biological pathways and molecular networks associated with variability in Ab responses. Thus, IPA was performed on 46 annotated DE genes (out of 65) identified in the pre-vaccination blood of pigs classified as high or low responders based on IAV-specific IgG levels at 21 dpv. Twenty-seven biological functions or diseases were significantly enriched (p <1 × 10; Supplementary Table 10). Among these, three functions (“migration of cells”, “leukocyte migration”, and “maturation of blood cells”) were predicted to be decreased in low responders (z-score <–2), and several additional functions associated with cell movement and immune trafficking showed similar trends. Moreover, inflammatory and immune-related processes were also predominantly predicted to be decreased in low responders, whereas functions related to cell death and survival tended to be increased.

“Immunoglobulin” was identified as the top upstream regulator (p = 1.65 × 10). Network analysis revealed five molecular interaction networks (Supplementary Table 11). The highest-scoring network (score = 41) was linked to “hematological disease, immunological disease, organismal injury, and abnormalities” and was centered around immunoglobulins (Fig. 4). Among the genes present in this network, some DE genes were associated with cell migration (*NRP1, NTN4, VNN2, NLRP3,* and *TYROBP*), inflammatory and immune responses (*ALOX15, ALOX5AP, IL1A, LGALS3, CEBPD, CRISPLD2, NFIL3,* and *CFD*), signaling and immune regulation (*BMX, RAPGEF4, TNFSF13B, DUSP1, DUSP2,* and *GAPT*), stress, apoptosis, or metabolic pathways (*ACSL4, AGTRAP, BNIP5,* and *MFAP3*). In this network, all genes except *NRP1* and *NTN4* were upregulated in low responders.

**Figure 4.**
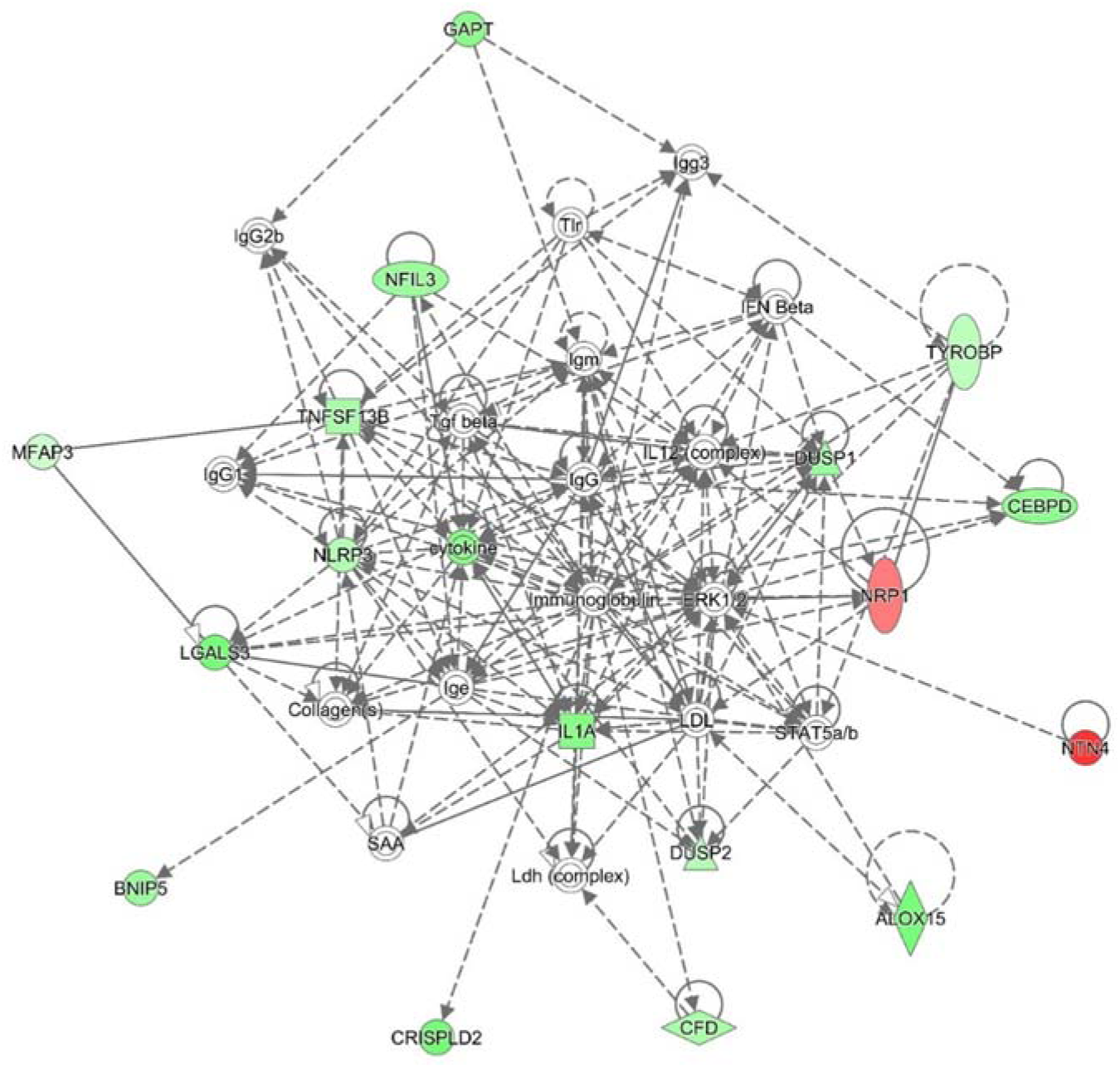
Gene interaction network of genes predisposing to an early antibody response to IAV vaccination. Network generated using Ingenuity Pathway Analysis (IPA), representing the top-ranked network associated with genes differentially expressed (DE) in the pre-vaccination blood transcriptome of animals exhibiting high or low serum IAV-specific IgG levels at 21 days post-vaccination (dpv). The network is mainly enriched for functions related to “hematological disease, immunological disease, organismal injury and abnormalities”. Genes upregulated in high responders are shown in red, while genes upregulated in low responders are shown in green

#### Functional analysis of pre-vaccination blood transcriptomic profiles between high and low responders at 118 dpv

IPA was performed on 186 annotated DE genes (out of 243) identified in the pre-vaccination blood of pigs classified as high or low responders based on HAI titers at 118 dpv. Ninety-one diseases or biological functions were significantly enriched (p < 1 × 10; Supplementary Table 12), the majority related to cancer, as a common bias of IPA analyses. Among the remaining functions, the function “organismal death” was predicted to be decreased in low responders (z-score = –3.195), whereas DNA repair processes (“double-stranded DNA break repair” and “DNA repair”) were predicted to be increased (z-scores = 1.518 and 2.012, respectively). Several cell-cycle-related functions were also enriched, although with non-significant or non-calculable z-scores.

Network analysis identified 11 molecular interaction networks (Supplementary Table 13). Network #3 (score = 52) was associated with “cell cycle, hematological system development and function, and humoral immune response” (Fig. 5a), while network #4 (score = 34) was linked to “cell-mediated immune response, hematological system development and function, and small molecule biochemistry”, and also included immunoglobulins (Fig. 5b). Representative genes in these networks included *FOXM1, KNTC1, MELK, POLA1, POLQ, STIL, LIN54, MED12L,* and *MED27* (cell cycle and DNA repair); *AFF4, REL, PKN3, PAG1, ITCH, RAPGEF4, MINDY3, TESK2,* and *ZNF250* (signal transduction and transcriptional regulation); *CD6, ITGAV, SKAP1, ITPKB, ST8SIA1, UBE2J2, ABCB10, ABCG1, NPRL3,* and *ZEB1* (immune and hematological functions); and genes such as *ISCU, FECH, GSPT1, PPME1, APBA3,* and *ARL15* involved in metabolic and mitochondrial processes. Other DE genes, including *EDC3*, *GIGYF1*, *GRIPAP1*, *CSPP1*, *SNX4*, and *REXO5*, were associated with RNA metabolism and protein turnover.

**Figure 5.**
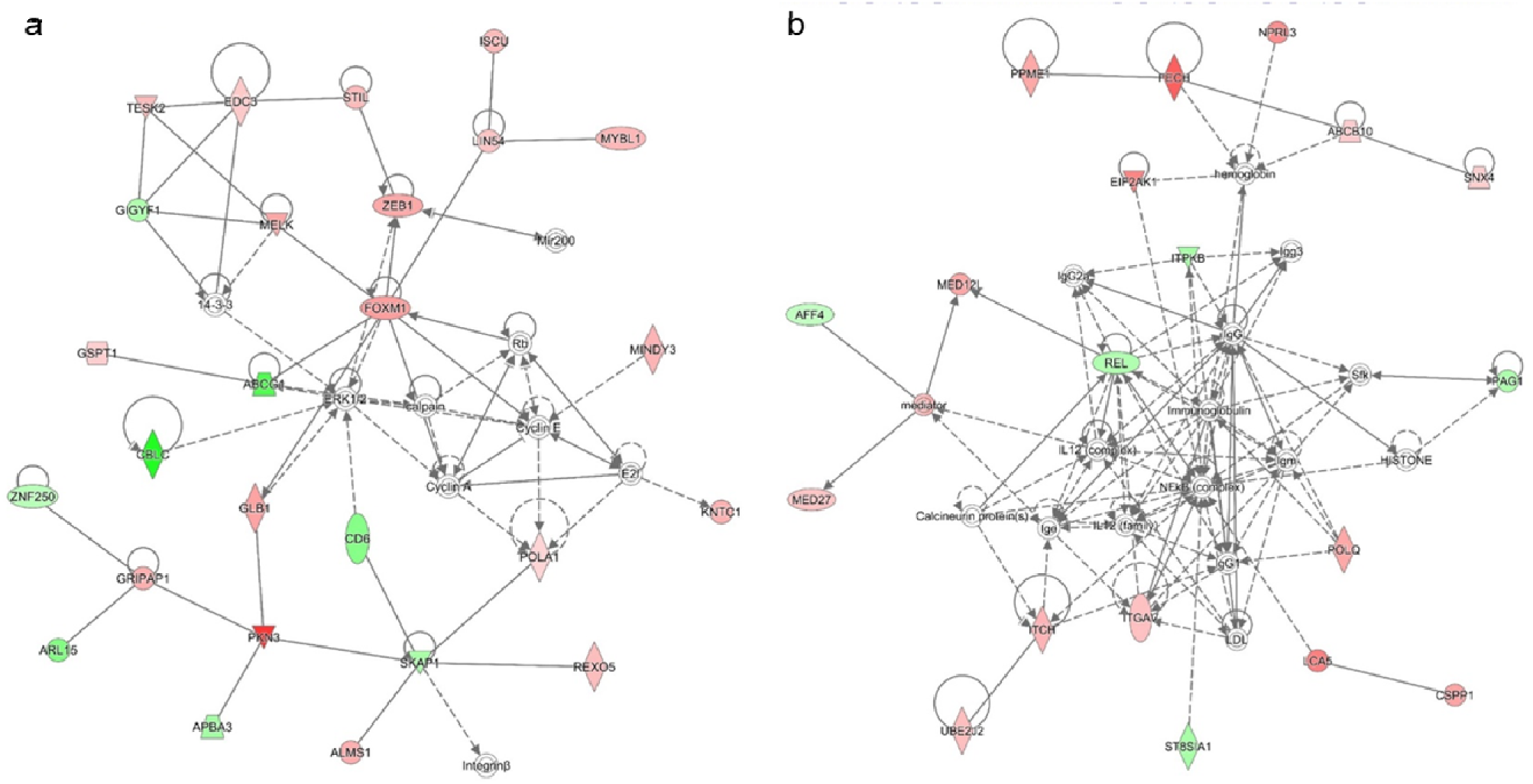
Gene interaction networks of genes predisposing to a persistence of the antibody response to IAV vaccination. Networks generated using Ingenuity Pathway Analysis (IPA), representing two top-ranked networks associated with genes differentially expressed (DE) in the pre-vaccination blood transcriptome of animals exhibiting high or low HAI titers at 118 days post-vaccination (dpv). Genes upregulated in high responders are shown in red, while genes upregulated in low responders are shown in green. (a) Network #3: associated with “cell cycle, hematological system development and function, and humoral immune response”. (b) Network #4: associated with “cell-mediated immune response, hematological system development and function, and small molecule biochemistry”.

#### Feature set enrichment analysis (FSEA) of pre-vaccination blood transcriptomes among animals with contrasted antibody responses

Given the limited number of DE genes at certain time points, FSEA was applied to ranked gene lists (based on absolute logFC) from all high versus low responder comparisons (Fig. 6; Supplementary Table 14). This approach enabled the detection of broader transcriptomic signatures associated with the intensity of the antibody response.

**Figure 6.**
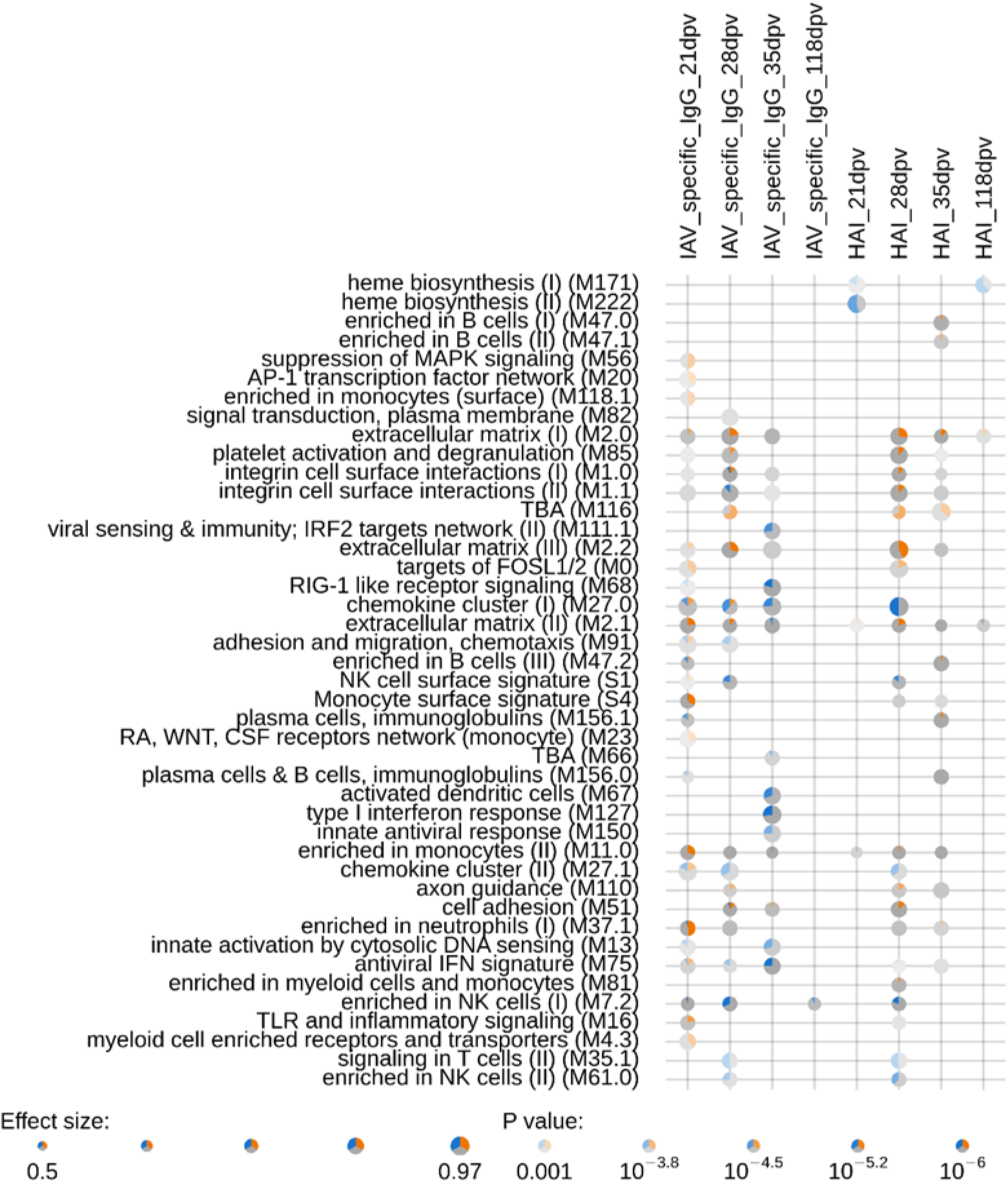
FSEA of genes differentially expressed between groups of animals with contrasted responses to IAV vaccination. Functional Set Enrichment Analysis (FSEA) performed using the CERNO test on gene lists ranked by the absolute log fold change (logFC) from differential expression (DE) analyses comparing animals with high versus low responses to vaccination at 21, 28, 35, and 118 days post-vaccination (dpv). Blood Transcriptional Modules (BTMs) significantly enriched in each list (*p* <0.001 and AUC >0.75) are shown in rows. Each BTM is labeled by its title and ID (in parentheses). The color transparency reflects the strength of the *p*-value, while the plot size represents the effect size (AUC). Significantly upregulated genes (*p* <0.05) in “high” responders are shown in blue, and downregulated genes in orange; non-significant genes are shown in grey. TBA = to be annotated.

For IAV-specific IgG levels, at 21 dpv, low antibody responders showed enrichment for antiviral sensing pathways, including “innate activation by cytosolic DNA sensing” (M13) and “RIG-I-like receptor signaling” (M68). In contrast, high responders displayed enrichment for extracellular matrix (ECM) modules (M2.0, M2.1, M2.2), inflammatory signatures (M0, M16), and myeloid-related modules (M4.3, M11.0, M23, M37.1, M118.1, S4). At 28 dpv, low responders were enriched in terms of modules associated with cell migration (M91), inflammation (M27.0, M27.1), NK cell signature (M7.2, M61.0, S1), and T cell signaling (M35.1). At 35 dpv, antiviral (M13, M68, M75, M111.1, M127, M150), dendritic cell/antigen presentation (M67), and inflammatory (M27.0) modules remained enriched in low responders, whereas high responders displayed ECM/adhesion (M1.0, M1.1, M51, M2.0, M2.1, M2.2), and platelet activation (M85) signatures. At 118 dpv, only a few modules were enriched, including the NK cell module (M7.2) in low responders and the ECM modules (M2.0, M2.1) in high responders.

Overall, similar enrichment patterns were observed for HAI titers and IAV-specific IgG levels. Differences were mainly observed with respect to heme biosynthesis modules (M171, M222) and a subset of B cell-related modules (M47.0, M47.1), which were only found for HAI titers, as well as innate and anti-viral response modules (M67, M127, M150), which were only found for IAV-specific IgG levels.

### Identification of biomarkers in pre-vaccination blood predicting high and low antibody responses to the IAV vaccine

Because HAI titers reflect viral strain-specific neutralizing antibody responses, we focused our biomarker discovery on this trait rather than on total IAV-specific IgG levels. Sparse Partial Least Squares Discriminant Analysis (sPLS-DA) was carried out to identify predictive genes distinguishing between high and low HAI responders at 21, 28, 35, and 118 dpv (Supplementary Table 15). The resulting gene signatures and their loadings are listed in Supplementary Table 16. Notably, *MYL4* and *SMIM12* were consistently selected at both 35 and 118 dpv, suggesting a stable predictive role across the peak and persistent phases of the antibody response.

Predictive genes identified across all time points were combined into a set of 102 genes. This panel was evaluated using PLS-DA to classify animals as high or low responders for HAI titers (Fig. 7; Supplementary Table 17). For HAI titers, the first PLS-DA component alone achieved excellent accuracy (AUC >0.89), and near-perfect classification was obtained with two or three components (AUC >0.99). Significant balanced error rates (BER) were observed for all time points, ranging from 0.21 to 0.24 at 21, 28, and 35 dpv. BER were minimal at 118 dpv (0.08, 0.05, and 0.03 with one, two, or three components, respectively). The same 102-gene panel also accurately predicted high and low IAV-specific IgG responders (AUC >0.87), confirming the robustness of this predictive signature across various antibody traits and time points, including long-term persistent immunity responses at 118 dpv (Fig. 7; Supplementary Table 17).

**Figure 7.**
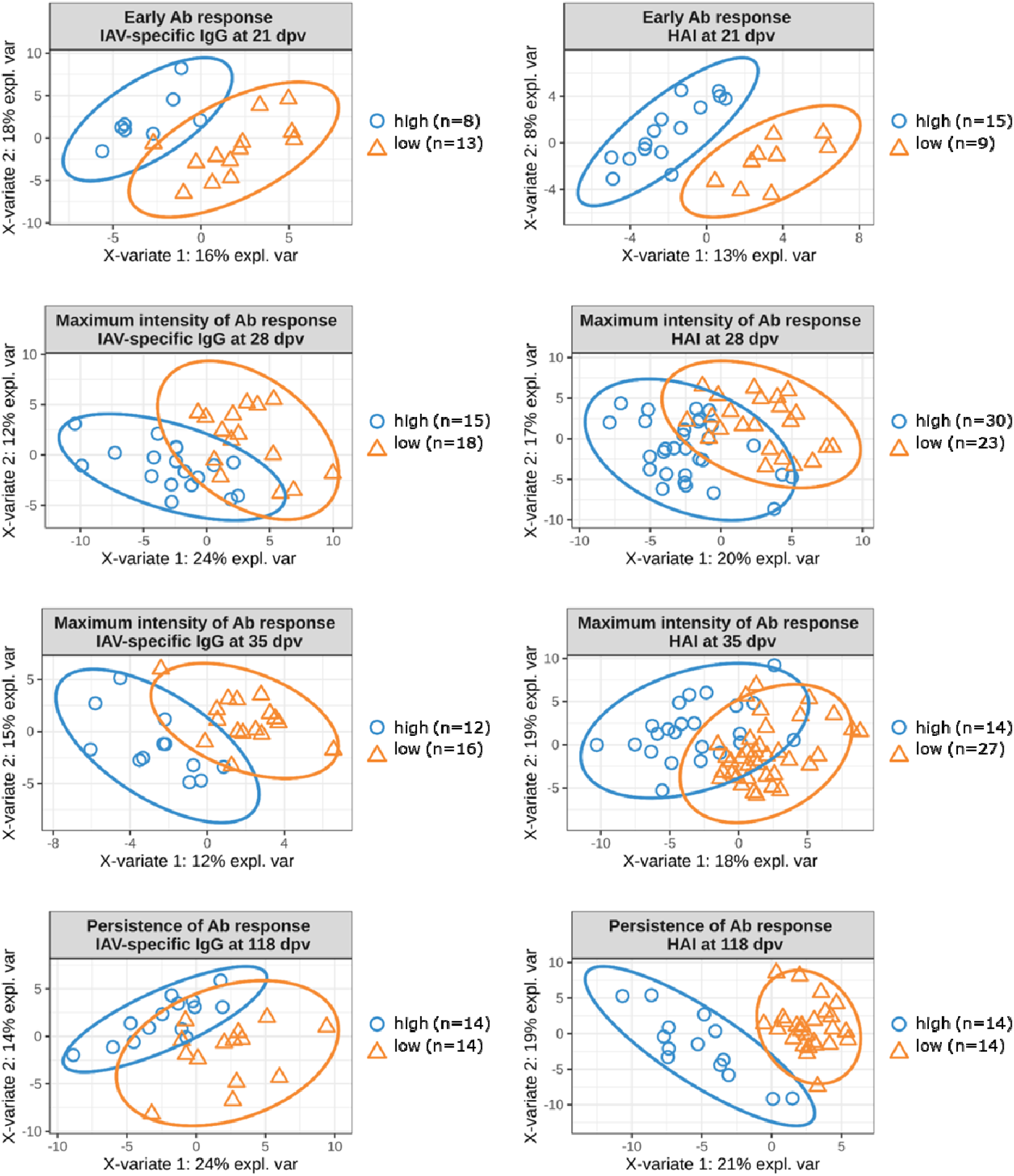
PLS-DA analysis based on 102 candidate predictive genes for the IAV vaccination response. Projection of groups of extreme responders (“high” vs “low”) for IAV-specific IgG or HAI titers at 21, 28, 35, and 118 days post-vaccination (dpv) into the subspace defined by the first two components of a Partial Least Squares Discriminant Analysis (PLS-DA) performed with the 102 candidate predictive genes. Confidence ellipses for each class are shown, with the confidence level set at 95%.

Among the 102 predictive genes, 14 corresponded to DE genes associated with HAI titers at 118 dpv (*FECH*, *TTC39B*, *PLGRKT*, *CST6*, *ENSSSCG00000014173*, *SEPT8*, *ENSSSCG00000034019*, *NLRP3*, *ENSSSCG00000037307*, *ENSSSCG00000038646*, *COX17*, *ITSN1*, *CFAP58*, and *UBAC1*). Another three genes (*C3orf52*, *DNAH9*, and *MYL4*) have previously been reported to be genetically regulated for transcription in pig blood transcriptome at 60 days of age ^34^.

Taken together, these findings identified a robust pre-vaccination transcriptional signature that could predict both the magnitude and persistence of the antibody response to IAV vaccination in pigs, highlighting potential biomarkers of vaccine responsiveness.

## Discussion

During this study, we observed a strong variability among pigs in terms of their antibody responses to IAV vaccination, despite the fact that all the animals had been vaccinated under identical conditions and produced specific antibodies. Nearly all piglets achieved protective HAI titers after the booster; however, the persistence of antibodies varied among individuals. Persistence at 118 dpv proved to be the most informative trait, supported by two genome-wide association signals and by the strongest transcriptomic predictors. Interestingly, blood gene expression before vaccination already reflected future antibody performance, in both the short and longer terms. This aligns with findings in humans, where baseline immune transcriptional states predict responses to influenza and other vaccines ^27–30^. We had previously revealed similar patterns in pigs vaccinated against *Mycoplasma hyopneumoniae* ^13^; here, we have extended this paradigm to influenza. Together, our observations support that individual variations in immune parameters before vaccination influence the magnitude and durability of antibody responses to vaccination in pigs.

### Influence of maternally-derived antibodies

Maternal antibodies can reduce the likelihood of seroconversion in piglets ^35–37^. Here, we did not detect an effect on HAI titers, likely because levels of maternal neutralizing antibodies were either low or undetectable at the time of vaccination. However, maternal IgG could not be distinguished from vaccine-induced IgG, so IgG analyses were restricted to MDA^-^ piglets. This improvement in accuracy came at the cost of a reduced animal sample size, particularly for transcriptomic and GWAS analyses, which consequently limited statistical power.

### Limited but significant trade-off between immune response magnitude and animal growth traits

Pigs with the strongest responses at 35 dpv displayed slightly lower growth performance (around 4–7% lower final weight and ∼5% lower ADG). While modest, such differences matter in production systems. Similar immunity-growth trade-offs have been reported elsewhere ^13,38^. If breeding programs prioritize growth over immune traits, these trade-offs may become more pronounced. This supports the inclusion of immune resilience traits in breeding goals. From an ecological immunology perspective ^39^, very strong responses are not always optimal: once protection is achieved, additional antibody production may not offer any benefits and can even reduce growth ^40^. In livestock systems, achieving sufficient rather than maximal immunity may represent a more balanced strategy, thereby minimizing the potential negative impacts of trade-offs between immunity and growth.

### Defining a “good” vaccine response: the importance of persistence

Assessing the performance of influenza vaccination requires immune readouts that reflect meaningful protection against the disease. In humans, HAI titers remain the most widely used correlates, with the classical cut-off of ≥40 historically linked to ∼50% protection ^41^. This threshold depends on the animal population and viral strain and was initially established in humans ^42^. Although useful, HAI alone does not always predict protection, as individuals can still become infected despite hypothetically protective titers ^43^. Other immune components (including T-cell responses, IFN-γ-producing cells, neuraminidase-inhibiting antibodies, and mucosal IgA) also contribute to protection, but none serve as a universal correlate ^44^.

A similar situation exists in pigs. Several immune readouts have been characterized ^45^, but no single gold-standard correlate of protection has been established. HAI ≥40 is nevertheless commonly used in swine practice ^46^, while ELISA tests are primarily employed for surveillance rather than predicting protection^47^.

In our cohort, nearly all the pigs exceeded the HAI threshold after boosting (28–35 days post-vaccination), indicating that early titers did not help to discriminate between animals. Instead, persistence at 118 dpv (covering most of the production lifespan) indicated pronounced variability and proved to be the most informative trait. Importantly, this late time point coincided with the strongest GWAS signals and the highest predictive accuracy from baseline transcriptional models. This suggests that persistence, rather than peak titers, is a key indicator of vaccine-induced humoral immunity in pigs.

### Distinct biological programs over the three phases of the vaccine response

Our data indicate that different biological processes drive early priming, post-boost amplification, and long-term antibody persistence. Piglets that responded rapidly after the first dose showed baseline enrichment for ECM remodeling, cell-adhesion pathways, and myeloid/dendritic readiness, features likely to support efficient antigen capture and the initiation of germinal center activity. By contrast, low early responders displayed stronger type I interferon and NK-cell transcriptional tone. After boosting, transcriptional differences between high and low responders decreased markedly, suggesting that the second dose largely overcame baseline variability and elicited robust responses in most animals. However, at 118 dpv, the early-response pattern re-emerged: ECM- and myeloid-associated signatures predicted sustained antibody levels, while interferon-related signatures were linked to poorer persistence. These observations support a model in which the “foundations” for durable humoral immunity are laid before vaccination and reflected in early priming, while peak titers depend more directly on vaccine stimulation.

This pattern mirrors findings from studies on *Mycoplasma hyopneumoniae* vaccination in pigs ^13^, and on human influenza ^29^ and SARS-CoV-2 ^30^: individuals with high interferon/inflammatory baseline activity tended to respond less well, whereas myeloid and stromal readiness favored stronger, longer-lasting responses. Mechanistic work supports this framework: stromal and extracellular matrix programs have been shown to influence the structure of the germinal center and the survival of long-lived plasma cells ^48^. Conversely, sustained type I interferon activity can disrupt germinal center organization and ultimately weaken antibody responses ^49^.

It is notable that these patterns were very similar to those observed in our previous study on *Mycoplasma hyopneumoniae* vaccination ^13^. Two studies were conducted in parallel using sibling animals from the same litters, reared under identical conditions, and vaccinated against either *Mycoplasma hyopneumoniae* or IAV under comparable experimental designs. Despite the distinct nature of the pathogens and vaccines, similar baseline transcriptional programs were associated with vaccine responsiveness, including the enrichment of extracellular matrix, stromal, and myeloid-related modules in high responders, as well as interferon-and inflammatory-related signatures in low responders. This convergence across independent vaccination contexts supports the existence of shared, host-intrinsic immune programs that shape the magnitude and persistence of vaccine-induced antibody responses in pigs.

### Genetic control and convergent immunological pathways

Our GWAS identified two regions in SSC5 and SSC8 associated with persistence. These regions include genes involved in T-cell help (*IL2RB*, *CSF2RB*), NF-κB signaling (*CARD10*), B-cell activation and cytoskeletal remodeling (*RAC2*), Notch-mediated lymphocyte differentiation and persistence (*RBPJ*), and Ca² homeostasis, which plays a role in long-term memory formation (*STIM2*). Importantly, several genes located in these GWAS regions were also differentially expressed before vaccination and featured in predictive models (e.g., *PLGRKT*, *CST6*, *ITSN1*, *TIMP3*, and *TBC1D19*), thus providing multiple independent lines of evidence for their involvement.

### Predictive value of pre-vaccination transcriptomic signatures

As well as identifying biological pathways associated with vaccine responsiveness, our study was also able to demonstrate that pre-vaccination blood transcriptomic profiles are of predictive value regarding both the magnitude and persistence of antibody responses to IAV vaccination. Using internally validated multivariate models, baseline gene expression signatures accurately discriminated between high and low responders across multiple time points, with a particularly strong performance for long-term persistence at 118 days post-vaccination (dpv). These findings thus support the concept that vaccine-induced humoral immunity is, at least in part, pre-configured by host-intrinsic immune states present prior to vaccination.

Importantly, these predictive signatures were consistent with the biological programs identified through differential expression and enrichment analyses, notably involving extracellular matrix remodeling, myeloid cell readiness, and interferon-related pathways. This concordance between mechanistic insights and predictive modeling reinforces the biological relevance of the signatures identified. While these models were derived and validated internally and will require confirmation in independent cohorts, they highlight the potential of baseline blood transcriptomics as a tool to stratify animals according to their expected vaccine responsiveness. In the context of livestock production, such approaches could support precision vaccination strategies and inform breeding programs intended to improve immune competence and vaccine efficiency.

### Beyond genetics and blood transcriptomics, additional determinants of vaccine responsiveness

Together, these findings highlight the contribution of host genetics and pre-existing immune transcriptional states to vaccine responsiveness. However, vaccine-induced immunity is a multifactorial trait, and other biological layers may further contribute to inter-individual variability.

In addition to genetics and blood transcriptomics, other layers of biological information—such as proteomics, glycomics, and metabolomics—can also contribute to determining inter-individual variability in vaccine responses ^50^. The microbiota has emerged as a potential source of biomarkers that influence vaccine efficacy. We previously conducted studies investigating the relationships between fecal microbiota composition and immune responses to *Mycoplasma hyopneumoniae* and IAV vaccination in pigs ^51,52^, and showed that specific pre-vaccination gut microbial profiles were associated with, and in some cases predictive of, the magnitude of antibody responses following vaccination.

## Conclusion

Both genetics and pre-existing immune status shape IAV vaccine responses in pigs. Among the traits measured during this study, antibody persistence proved to be particularly informative and biologically meaningful. Our findings support the concept that persistent humoral immunity is not solely a downstream result of vaccination, but rather reflects the inherent immune programs present prior to immunization. Our results therefore offer a means to develop biomarkers predictive of vaccine response magnitude. A clearer understanding of the determinants of immune competence could be expected to inform precision vaccination strategies and improve breeding programs, leading to greater sustainability and disease resistance.

## Methods

### Animal design, production phenotypes and sampling

This study involved 48 families of Large White pigs that were produced in five batches and raised without antibiotic treatment at the GenESI, INRAE, Pig Innovative Breeding Experimental Facility, https://doi.org/10.15454/1.5572415481185847E12. Sows (n = 47) were inseminated with boar semen (n = 48) chosen to maximize genetic variability. Within each litter, piglets of each sex were selected based on their BW at 21d, the aim being to represent the litter average rather than favor the heaviest or smallest animals. In total, 276 piglets (143 uncastrated males and 133 females) were enrolled. Three to five piglets per litter received the commercial influenza A vaccine (Respiporc Flu3, IDT Biologika), while one or two littermates remained unvaccinated (in total, 201 vaccinated and 75 non-vaccinated animals). The first dose was administered at weaning (0 dpv, mean age 28 days; range 24–31 days), with a booster at 21 dpv (Fig. 1a). A small number of animals was removed from the study for reasons unrelated to vaccination, including farming-related health issues leading to morbidity or premature death, sampling errors, or, in one case, an ambiguous lack of response to vaccination where we could not conclude whether it was a non-responder or if the vaccination injection had failed. The final dataset consisted of 187 vaccinated and 63 control piglets (Supplementary Table 1), of which 173 and 60, respectively, were monitored until 118 dpv. Housing conditions followed standard pig production practices, with animals grouped in pens of 20–30 pigs during post-weaning (from 28 to 68 days) and 10–12 pigs during the finishing period (from 68 to 146 days), and all receiving the same commercial diet.

BW was recorded at birth, shortly before weaning (around 21 days), at weaning (0 dpv), at the end of the post-weaning phase (40 dpv), and again prior to slaughter (118 dpv). ADG was calculated for the periods 0–40 dpv and 40–118 dpv. Blood was sampled from the jugular vein at 0, 21, 28, 35 and 118 dpv for serum preparation, and at 0 dpv for DNA and RNA extraction. Blood was collected in dry tubes for serum preparation, in EDTA tubes (Becton Dickinson) for DNA study and in Tempus stabilizing tubes (Thermo Fisher) for RNA extraction. The samples were stored at−20°C (DNA) or −80°C (RNA) until they were extracted for analysis. Blood was also collected from the sows during the week preceding farrowing, in order to assess maternal antibody status.

### Measurement of IAV-specific IgG levels and HAI titers and classification of animals as high or low vaccine responders

The three influenza A strains used in this study—avian-like swine H1avN1 (A/Sw/Côtes d’Armor/0388/09), human-like reassortant swine H3N2 (A/Sw/Spain/SF32071/07) and human-like reassortant swine H1huN2 (A/Sw/Spain/SF12091/07) were kindly provided by Gaëlle Simon (ANSES, Maisons-Alfort, France). The viruses were propagated on MDCK cells for 72 h at a multiplicity of infection of 0.001. Cell supernatants were clarified by centrifugation (3600 rpm, 15 min, 4°C), inactivated with 0.037% formalin for 24 h at 37°C, and dialyzed against PBS to remove any residual formalin.

High-binding 96-well plates (NUNC Maxisorp) were coated overnight at room temperature with a mixture of the three inactivated strains (10 pfu/mL in PBS). After washing with PBS-0.05% Tween 20 and blocking with PBS-1% BSA, serum samples were diluted 1:100, 1:1000 or 1:10000 (in duplicate) and incubated for 90 min at room temperature. A standard curve was generated in parallel using anti-pig IgG-coated wells (A100-104A, Bethyl Laboratories, 1µg in 100µL/well) and a calibrated pig reference serum (RS10-107, Bethyl Laboratories, serial two-fold dilutions from 500 ng/mL). The plates were then incubated with HRP-conjugated anti-pig IgG (A100-104P, Bethyl Laboratories, 1:40,000) for 90 min, washed, and developed for 20 min (R&D Systems substrate reagent), followed by a reaction stop with 2N H□ SO□. Absorbance was read at 450 nm, and IgG concentrations were calculated from the standard curve and expressed in µg/mL.

HAI titers were determined as previously described ^53^. Sera were treated with a receptor-destroying enzyme (RDE; Denka Seiken) at 37°C overnight (1:4 dilution), heat-inactivated at 56°C for 30 min, and further diluted to 1:10 in PBS. The three inactivated strains were titrated before each assay to confirm the number of hemagglutination units, and then mixed to obtain 4 HA units. Two-fold serial serum dilutions starting at 1:10 were incubated with the virus for 30 min at room temperature, followed by the addition of turkey red blood cells (Tebu-Bio, 0.5%) and a further 30-min incubation. Titers were recorded as the reciprocal of the highest dilution preventing hemagglutination. Titers below detection (<5) were assigned a value of 2.5. Under these conditions, seroprotection corresponded to an HAI level of 40 or higher.

For IAV-specific IgG, extreme responders were identified among the piglets that lacked maternally derived antibodies at the time of vaccination. IAV-specific IgG values (log - transformed) measured at 21, 28, 35 and 118 dpv were used to define high and low responders as animals exceeding the mean +1 SD or below the mean –1 SD at each time point. For HAI titers, high and low responders were defined based on population distributions at each time point: ≤ 20 vs ≥ 160 (21 dpv), ≤ 160 vs ≥ 640 (28 and 35 dpv), and ≤ 10 vs ≥ 80 (118 dpv). Group compositions and descriptive statistics are provided in Supplementary Tables 1 and S2.

### Effects of zootechnical parameters on vaccine response and of vaccination on growth traits

All statistical analyses were performed using R (version 3.6.1). Antibody measurements were transformed prior to analysis (log for IAV-specific IgG concentrations and log for HAI titers) to meet model assumptions. To assess the effect of management-related factors on antibody responses, we fitted linear mixed-effects models using the lmer function from the lme4 package, including sex and batch as fixed effects, age at weaning (24–31 days) as a linear covariate, and litter as a random effect. The same framework was used to evaluate the impact of vaccination on growth traits, with vaccination status (vaccinated vs. non-vaccinated) or responder status (high vs. low responders) included as additional fixed effects. The significance of fixed effects was tested using the lmerTest, based on type III ANOVA with Satterthwaite’s approximation for degrees of freedom, and random-effect significance was assessed by likelihood-ratio testing. Post-hoc pairwise comparisons among batches were performed with the Tukey adjustment using emmeans. Correlations between immune and growth traits were explored using Pearson correlation matrices (corrplot). A significance threshold of 0.05 was applied throughout.

### Genome-wide association studies (GWAS)

Genomic data were generated for the vaccinated pigs (n = 187) using the Affymetrix Axiom Pig HD array (658K SNPs). Raw genotypes underwent standard quality control with the Axiom Suite, including Dish QC, sample call-rate and plate-level metrics. Only annotated autosomal markers were retained (598,138 SNPs; Axiom_PigHD_v1.na35.r4.a2.annot.csv). Further filtering was performed in R using the check.marker function from the GenABEL package, removing variants with a minor allele frequency below 5%, a call rate below 95%, or significant deviation from the Hardy-Weinberg equilibrium (FDR < 0.1). After filtering, 425,689 SNPs were retained for all 187 individuals. No outlier samples were detected, and genomic kinship estimates were consistent with pedigree records.

GWAS were conducted with RepeatABEL, fitting a linear mixed model that included batch and sex as fixed effects, age at weaning as a covariate, and random effects accounting for litter and the genomic relationship matrix. Associations were declared significant at FDR <0.05. Candidate genomic regions were mapped to the Sscrofa11.1 reference genome (Ensembl release 103).

### Blood transcriptome before vaccination

Whole-blood RNA was extracted from 92 vaccinated piglets without maternally-derived antibodies (Supplementary Table 1) using the Norgen Preserved Blood RNA Purification Kit I adapted for Tempus-stabilized samples. RNA quantity and integrity were assessed with a NanoDrop 2000 and Agilent Bioanalyzer, respectively (mean yield 100.2 ± 27.0 µg; mean RIN 7.8 ± 0.5, range 6.5–9). Libraries were generated from 1 µg total RNA using the Illumina TruSeq Stranded Total RNA kit with Ribo-Zero Globin depletion following the manufacturer’s instructions. After rRNA and globin removal, RNA was enzymatically fragmented, converted to cDNA, adapter-ligated and amplified. Library quality was confirmed on an Agilent TapeStation and quantified by Qubit prior to 12-plex pooling. Sequencing was carried out on an Illumina HiSeq 3000 (150 bp paired-end), with each pool run across two lanes.

Reads were aligned to the Sscrofa11.1 reference genome (Ensembl v90) using TopHat (version 2.1.0), and gene-level counts were obtained with htseq-count (version 0.6.1.p1). Mean sequencing depth exceeded 60 million reads per sample; one sample with <20 million reads was excluded. The final dataset comprised 91 piglets spanning the phenotypic extremes for antibody responses (IAV-specific IgG and HAI titers across 21, 28, 35 and 118 dpv). These individuals reflected the broader vaccinated population, with comparable mean antibody levels (Supplementary Table 2).

### RNA-seq differential expression and functional enrichment analyses

Differential expression analyses were performed in R using edgeR (version 3.26.6). Raw gene counts were normalized with calcNormFactors and variance-stabilized using limma-voom (version 3.40.6). Linear models included sex and batch as fixed effects and age at weaning as a covariate. Likelihood-ratio tests were used to compare extremely high and low antibody responders at each time point and for each serological trait. *P*-values were adjusted for multiple testing using the false discovery rate (FDR), with significance defined at FDR <0.05.

Differentially expressed genes were interrogated with IPA (Qiagen; version 60467501) to identify enriched canonical pathways, upstream regulators, predicted activation states, and regulatory networks. Significance thresholds followed IPA recommendations (disease/function *p*-value <1×10 and absolute activation z-score >2). Consistency scores were used to support predictions of regulatory networks.

FSEA was performed using the tmod R package (version 0.40) ^54^, employing the CERNO method to assess enrichment among genes ranked by absolute log-fold change. Gene sets corresponded to Blood Transcriptomic Modules (BTMs) ^26^ mapped to the pig genome and described elsewhere ^55^. Enrichment results were visualized using tmodPanelPlot.

### Predictive modelling (sPLS-DA and PLS-DA)

Sparse partial least squares-discriminant analysis (sPLS-DA) was conducted with mixOmics (version 6.10.8) to identify pre-vaccination genes discriminating high versus low responders ^56^. Count data were preprocessed as for differential expression. Model tuning employed stratified repeated 5-fold cross-validation (100 iterations), initially exploring up to three components and 5–95 genes per component, and then refining the parameters within optimal ranges (up to 30 genes per component). Model performance was primarily assessed using the balanced error rate (BER). The final models retained one or two components, selecting the smallest number of genes that achieved optimal classification performance across time points. The union of predictors selected across time points yielded a set of 102 genes, which should be considered as an internally derived candidate signature. The predictive accuracy of this gene set was evaluated using PLS-DA with cross-validation for both HAI titers and IAV-specific IgG responses at 21, 28, 35 and 118 dpv.

## Data availability

The blood transcriptome (RNA-Seq) and genotype datasets generated and analyzed during this study are available from the corresponding author upon reasonable request. All other data generated or analyzed during this study are included in this published article and its Supplementary Information files.

## Acknowledgements

The authors would like to thank Isabelle Schwartz-Cornil, who coordinated the H2020 SAPHIR project that gathered a large, multidisciplinary consortium and provided us with the opportunity to design ad hoc experiments to study the individual variability of vaccine responses. We offer special warm thanks to Chiara Ferrandi (†) and Alessandra Stella, who helped produce the libraries for RNA-Seq at Parco Technologico (PTP, Lodi, Italy). We thank the INRAE Gentyane Genomic Services platform (Clermont-Ferrand, France) for performing animal SNP genotyping. This work was performed in collaboration with the GeT core facility, Toulouse, France (GeT, https://doi.org/10.15454/1.5572370921303193E12). The GeT core facility is supported by France Génomique National infrastructure, funded as part of the “Investissement d’avenir” program managed by Agence Nationale pour la Recherche (contract ANR-10-INBS-09). We thank all members of the Genetics, Microbiota, Health team at the INRAE GABI unit, particularly Jean-Jacques Leplat and Nicolas Bruneau, for their assistance at all stages of the project, including animal sampling. We also thank the pig team at the GenESI unit for their assistance with animal housing and sampling throughout the experiment.

## Funding

This project has received funding from the European Union’s Horizon 2020 Programme for research, technological development, and demonstration under Grant Agreement No. 633184.

This publication reflects the views only of the authors, and not the European Commission (EC). The EC is not liable for any use that may be made of the information contained herein.

## Author Contributions

FB, JE, MHPvdL and CRG conceived and designed the experiment; LR and YB were in charge of pig production at the experimental farm; GL managed animal sampling and organized the biobanking of all samples; OB produced the RNA sequencing data; FB characterized the vaccine response; JE and TM performed bioinformatics analyses; FB and JE performed GWAS analyses and bio-statistical analyses; CRG supervised the project; FB and CRG interpreted all the data and wrote the paper. All authors read and approved the final manuscript.

