## Supplementary Figures 1 - 2 for "Host genetics and pre-vaccination blood transcriptome as determinants of vaccine-induced immunity to Influenza A virus in swine"

### Slide 1
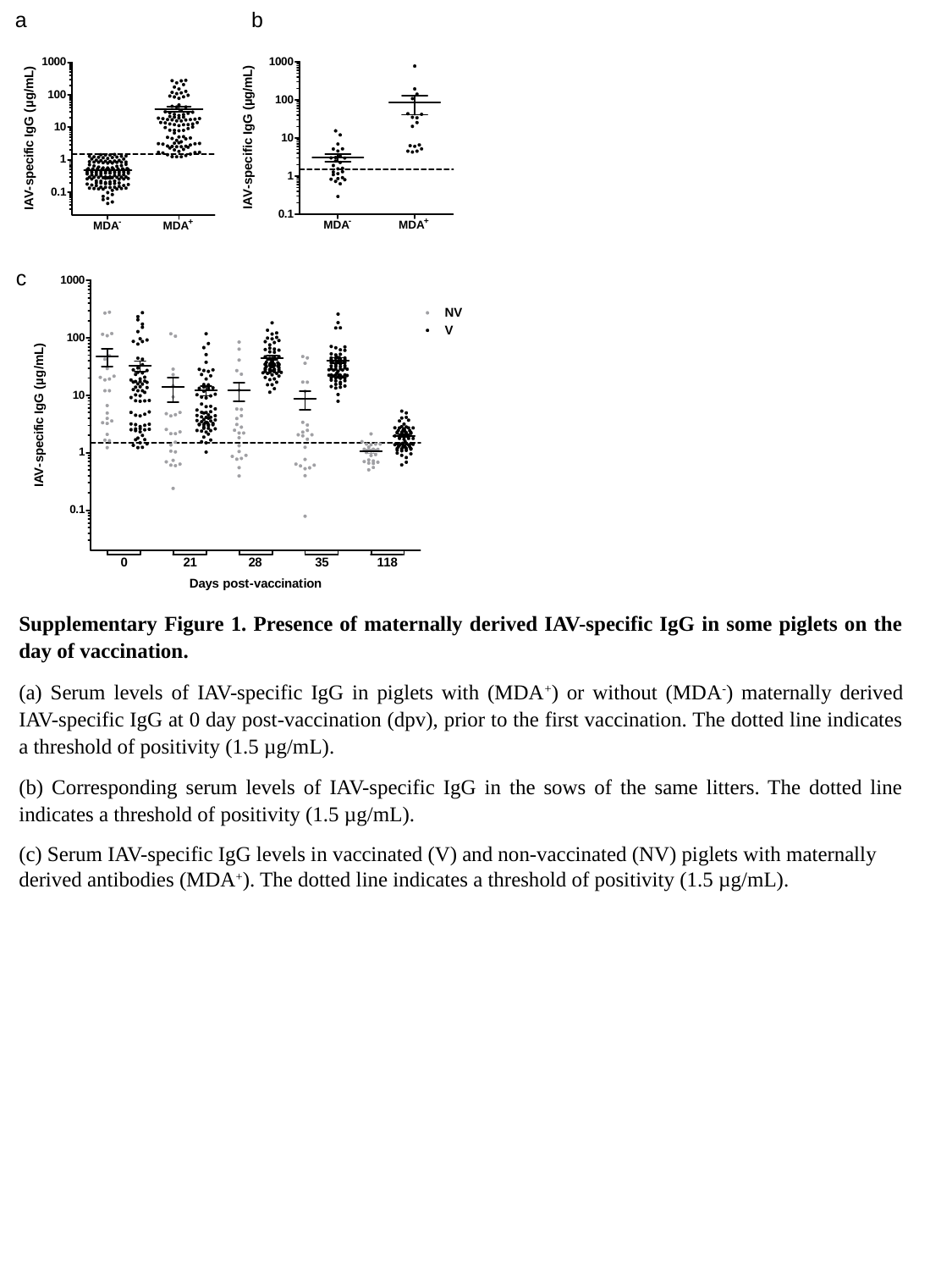

b
a
c
Supplementary Figure 1. Presence of maternally derived IAV-specific IgG in some piglets on the day of vaccination.
(a) Serum levels of IAV-specific IgG in piglets with (MDA^+^) or without (MDA^-^) maternally derived IAV-specific IgG at 0 day post-vaccination (dpv), prior to the first vaccination. The dotted line indicates a threshold of positivity (1.5 µg/mL).
(b) Corresponding serum levels of IAV-specific IgG in the sows of the same litters. The dotted line indicates a threshold of positivity (1.5 µg/mL).
(c) Serum IAV-specific IgG levels in vaccinated (V) and non-vaccinated (NV) piglets with maternally derived antibodies (MDA^+^). The dotted line indicates a threshold of positivity (1.5 µg/mL).

### Slide 2
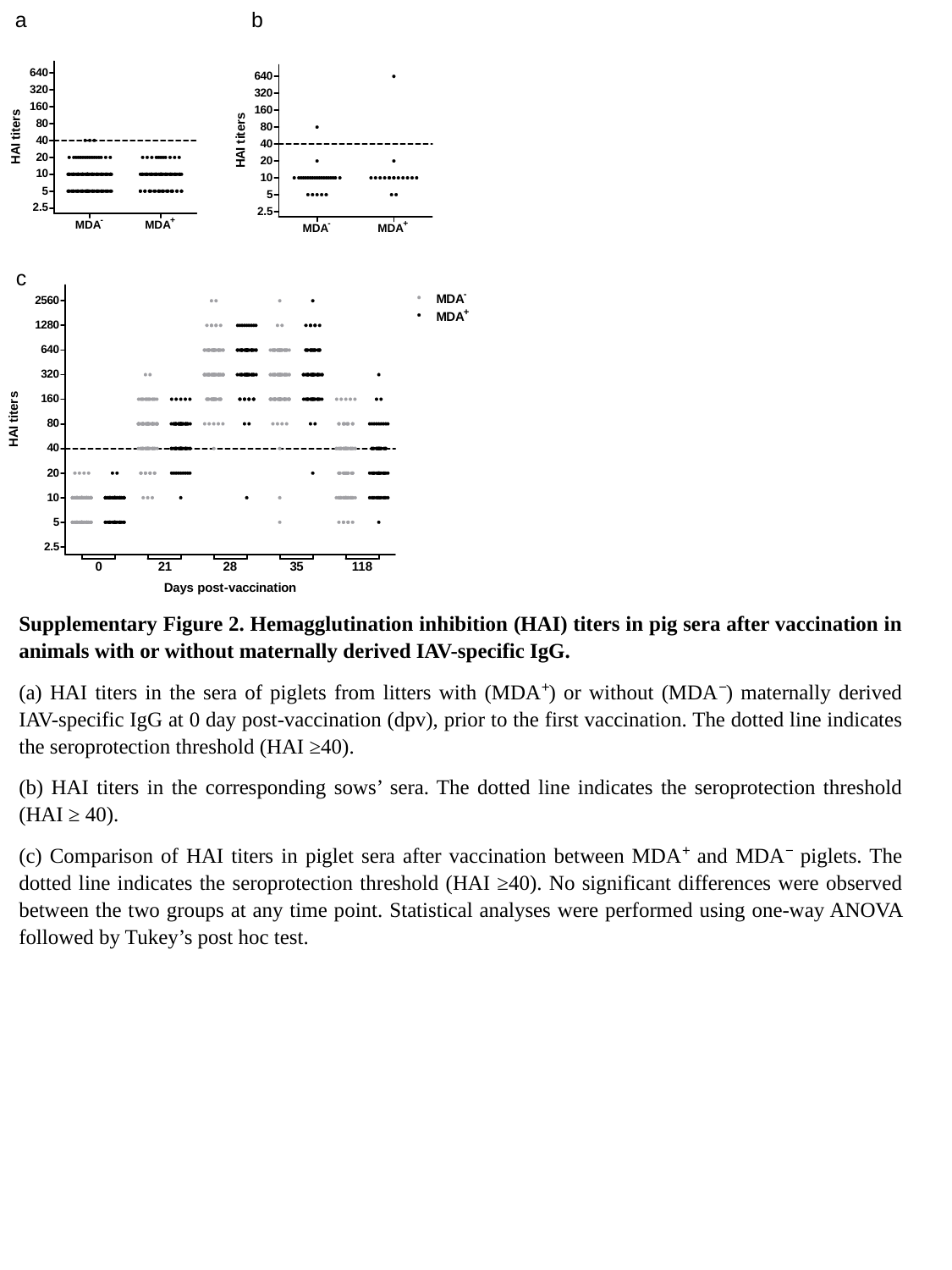

b
a
c
Supplementary Figure 2. Hemagglutination inhibition (HAI) titers in pig sera after vaccination in animals with or without maternally derived IAV-specific IgG.
(a) HAI titers in the sera of piglets from litters with (MDA⁺) or without (MDA⁻) maternally derived IAV-specific IgG at 0 day post-vaccination (dpv), prior to the first vaccination. The dotted line indicates the seroprotection threshold (HAI ≥40).
(b) HAI titers in the corresponding sows’ sera. The dotted line indicates the seroprotection threshold (HAI ≥ 40).
(c) Comparison of HAI titers in piglet sera after vaccination between MDA⁺ and MDA⁻ piglets. The dotted line indicates the seroprotection threshold (HAI ≥40). No significant differences were observed between the two groups at any time point. Statistical analyses were performed using one-way ANOVA followed by Tukey’s post hoc test.
